# Inference of Differential Gene Regulatory Networks Based on Gene Expression and Genetic Perturbation Data

**DOI:** 10.1101/466623

**Authors:** Xin Zhou, Xiaodong Cai

## Abstract

**Motivation:** Gene regulatory networks (GRNs) of the same organism can be different under different conditions, although the overall network structure may be similar. Understanding the difference in GRNs under different conditions is important to understand condition-specific gene regulation. When gene expression and other relevant data under two different conditions are available, they can be used by an existing network inference algorithm to estimate two GRNs separately, and then to identify the difference between the two GRNs. However, such an approach does not exploit the similarity in two GRNs, and may sacrifice inference accuracy.

**Results:** In this paper, we model GRNs with the structural equation model (SEM) that can integrate gene expression and genetic perturbation data, and develop an algorithm named fused sparse SEM (FSSEM), to jointly infer GRNs under two conditions, and then to identify difference of the two GRNs. Computer simulations demonstrate that the FSSEM algorithm outperforms the approach that estimates two GRNs separately. Analysis of a gene expression and SNP dataset of lung cancer and normal lung tissues with FSSEM inferred a GRN largely agree with the known lung GRN reported in the literature, and it identified a differential GRN, whose genes with largest degrees were reported to be implicated in lung cancer. The FSSEM algorithm provides a valuable tool for joint inference of two GRNs and identification of the differential GRN under two conditions.

**Availability:** The software package for the FSSEM algorithm is available at https://github.com/Ivis4ml/FSSEM.git

## Introduction

A gene regulatory networks (GRN) consists of a set of genes that interact with each other to govern their expression and molecular functions. For example, transcription factors (TFs) can bind to promoter regions of their target genes and regulate the expression of target genes (Harbison *et al*., 2004). Gene-gene interactions can change under different environments, in different tissue types or disease states, and during development and speciation (Ideker and Krogan, 2012). Therefore, GRNs undergo substantial rewiring depending on specific molecular context in which they operate (Califano, 2011). Identification of condition-specific GRNs is critical to unravel the molecular mechanism of various tissue or disease-specific biological processes (Sonawane *et al*., 2017). Although a number of computational methods have been developed to infer GRNs from gene expression and other relevant data, they are mainly concerned with the static structure of gene networks under a certain condition. Several methods aim to infer GRNs using only gene expression data; they include the approaches that construct relevance network based on a similarity measure, such as correlation or mutual information (Butte and Kohane, 1999; Faith *et al*., 2007; Margolin *et al*., 2006), Gaussian Graphical Model (GGM) (Friedman *et al*., 2008), Bayesian networks (Statnikov and Aliferis, 2010), and linear regression model (Haury *et al*., 2012). Several other methods infer GRNs by integrating genetic perturbations with gene expression data; these methods include approaches using Bayesian networks incorporating expression quantitative trait loci (eQTLs) (Zhu *et al*., 2007), likelihood-based causal models (Neto *et al*., 2008), and structural equation models (SEMs) (Cai *et al*., 2013; Liu *et al*., 2008; Logsdon and Mezey, 2010).

While it is possible to apply these methods to identify GRNs under different conditions separately, such an approach is apparently not optimal to identify the difference in GRNs, because it does not exploit the similarity in two GRNs. Several methods have been proposed to use the gene expression data of different conditions to jointly estimate GRNs under different conditions. Particularly, GRNs under multiple conditions are modeled with multiple GGMs, and these GGMs are inferred jointly from gene expression data (Danaher *et al*., 2014). When a gene is mutated, its regulatory effect on all its target gene may changed. Taking into account such effects, a node-based approach to joint inference of multiple GGMs were developed in (Mohan *et al*., 2014). GGMs exploit the sample covariance of the gene expression levels, but they cannot integrate genetic perturbations with gene expression data. Moreover, it has been demonstrated that genetic perturbation along with gene expression data can determine directed edges in GRNs (Logsdon and Mezey, 2010), but GGMs can only identify undirected edges.

In this paper, we employ SEMs to model GRNs as described in (Cai *et al*., 2013; Liu *et al*., 2008; Logsdon and Mezey, 2010). This enables us to integrate genetic perturbation data with gene expression data. Taking into account the sparsity in GRNs, we have developed a sparse-aware maximum likelihood (SML) method (Cai *et al*., 2013) to infer a single GRN based on SEM. Here, taking into account not only the sparsity in GRNs but also the sparsity in the differences between GRNs under two different conditions, we develop an algorithm, named fused sparse SEM (FSSEM), to infer two GRNs from different conditions jointly, and then to identify difference in two GRNs. Computer simulations demonstrate the superior performance of our novel approach relative to the existing one that infers GRNs under two conditions separately.

## Methods

### 1.1 GRN model

Suppose that expression levels of *n* genes under two different conditions are measured using e.g. micro-array or RNA-Seq technique. Let 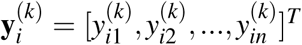 denote expression level of *n* genes in individual *i* under condition *k*, where *k* = 1,2 and *i* = 1,2,…, *n_k_*, with *n_k_* being the number of individuals where gene expression levels are measured under condition *k*. Supposed that a set of perturbations of these genes have also been measured. These perturbations can be due to e.g., eQTLs or gene copy number variants (CNVs). In this paper, we will consider only eQTLs. As in (Cai *et al*., 2013; Logsdon and Mezey, 2010), we assume that each gene in the GRN of interest has at least one cis-eQTL, so that the structure of underlying GRN is uniquely identifiable. Let 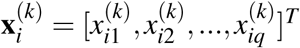 denote the genotypes of *q cis*-eQTLs in individual *i* under condition *k*, where *k* = 1,2 and *i* = 1,2,…,*n_k_*. Since the expression level of a particular gene may be regulated by other genes and is affected by its eQTLs, we employ the following SEM to model the expression of *n* genes

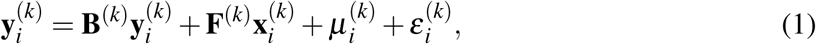

where *i* = 1,…, *n_k_*, *k* = 1,2, *n* × *n* matrix **B**^(*k*)^ defines the unknown network structure under condition *k*, *n* × *q* matrix **F**^(*k*)^ captures the effect of *cis*-eQTLs on gene expression levels under condition *k*, *n* × 1 vector ***μ***^(*k*)^ accounts for the model bias in SEM, and *n* × 1 vector ***ε***^(*k*)^ denotes the residual error, which is modeled as a Gaussian vector with zero mean and variance ***σ***^2^. It is assumed that no self-loops are presented per gene in GRN, which implies that the diagonal entries of **B**^(*k*)^ are zero, and it is also assumed that *q cis*-eQTLs have been identified using an existing eQTL method, but their regulatory effects are unknown, thus, **F**^(*k*)^ has *q* nonzero entries with known locations.

### 1.2 Joint Inference of two GRNs

Let 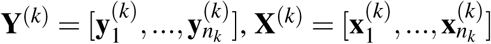 and 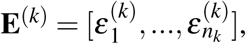 where *k* = 1,2, and assume that *n*_1_ + *n*_2_ observations are independent. Then, the negative log-likelihood function of the data can be written as

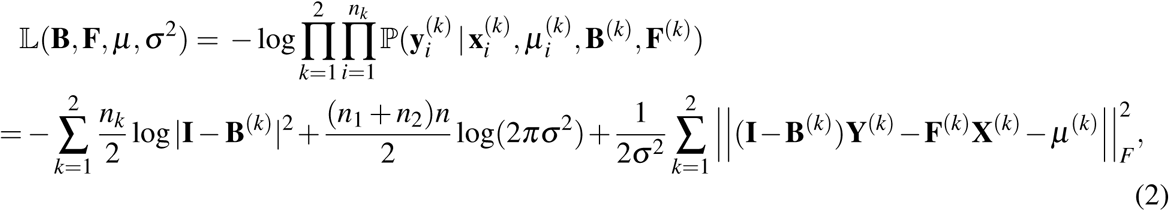

where **B** = [**B**^(1)^,**B**^(2)^], **F** = [**F**^(1)^,**F**^(2)^], ***μ*** = [***μ***^(1)^,***μ***^(2)^], and ∥ ⋅ ∥_*F*_ stands for the Frobenius norm. Our goal is to estimate **B**^(1)^ and **B**^(2)^ in (2). It is not difficult to show that minimizing (2) with respect to ***μ*** yields 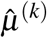 = (**I** − **B**^(*k*)^)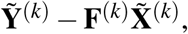 where 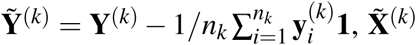 = 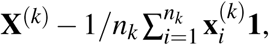 and **1** is a vector with all its entries equal to 1.

Since a gene is regulated by a small number of other genes (Gardner *et al*., 2003; Tegner *et al*., 2003; Thieffry *et al*., 1998), GRNs are sparse, meaning that most entries of **B**^(1)^ and **B**^(2)^ are zeroes. Moreover, it is reasonable to expect that changes in a GRN under two different conditions is relatively small. Therefore, most entries of **B**^(2)^ – **B**^(1)^ are zeroes. Let 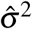 be an estimate of ***σ***^2^ that will be specified later, replacing ***μ***^(*k*)^ and ***σ***^2^ in (2) with 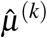 and 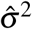 respectively, and taking into account the sparsity in **B**^(1)^ and **B**^(2)^, and the sparsity in **B**^(2)^ – **B**^(1)^, we can estimate **B** and **F** by minimizing the following penalized negative log-likelihood function

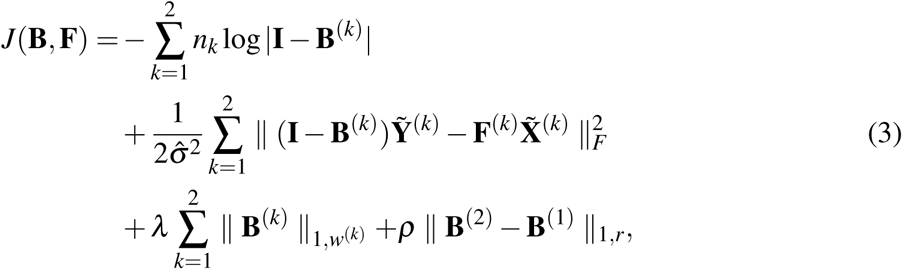

where 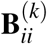 = 0, ∀*i* = 1,…, *n*, *k* = 1,2, ∥ **B**^(*k*)^ ∥_1,*w*(*k*)_ = 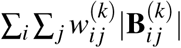 is the weighted *ℓ*_1_-norm, ∥ **B**^(2)^–**B**^(1)^ ∥_1,*r*_ is also a weighted *ℓ*_1_-norm with similar definition, ***λ*** and ***ρ*** are two nonnegative parameters. Weights 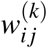 and *r_ij_* in the penalty terms are introduced to improve estimation accuracy and robustness in line with the adaptive lasso (Zou, 2006) and the adaptive generalized fused lasso (Viallon *et* al., 2016), and they are selected as 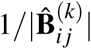 and 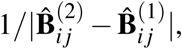 respectively, where 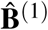 and 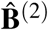 are preliminary estimates of **B**^(1)^ and **B**^(2)^ obtained from the following ridge regression:

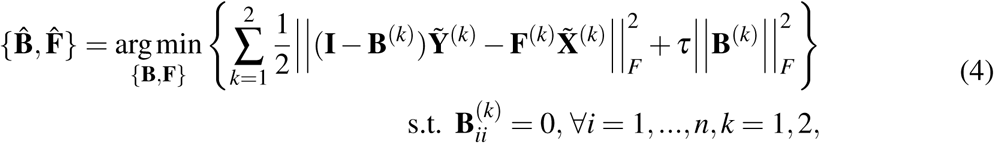

where 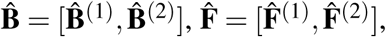 and the estimate of ***σ***^2^, 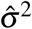 in (3), is given by

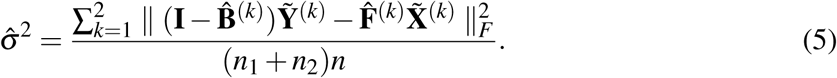

Based on (3), we next develop a proximal alternative linearize minimization algorithm to infer **B**^(1)^ and **B**^(2)^.

### 1.3 Ridge regression

In the first stage, we solve the ridge regression problem (4) to find initial values of **B**, **F**, weights *w*^(*k*)^, *k* = 1,2, and *r* for the FSSEM algorithm to minimize (3). Let 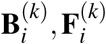 and 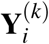 be the *i*-th row of **B**^(*k*)^, **F**^(*k*)^ and **Y**^(*k*)^, respectively. Define 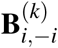 as the 1 × (*n* − 1) vector obtained by removing the *i*-th entry from 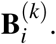 Let *S_q_*(*i*) be the set of indices of non-zero entries in the 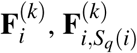 be the vector that contains the nonzero entries of 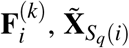 be the matrix formed by taking rows of 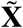 whose indices are in *S_q_*(*i*), and 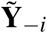 be the matrix formed by removing *i*th row of 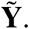

Then, the ridge regression problem (4) can be decomposed into *n* separate problems:

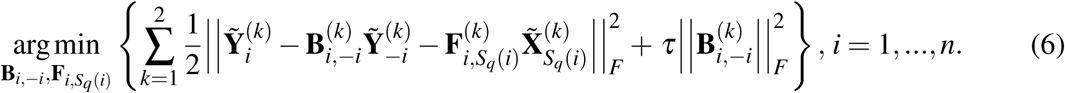

Minimizing the objective function in (6) with respect to (w.r.t.) 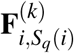 yields the following closed form solution

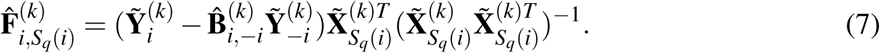

Substituting 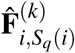 into (6) and minimizing w.r.t. 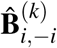 gives 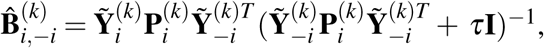 which in turn results in 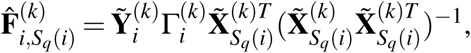 where 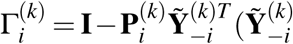 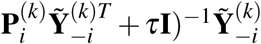 and 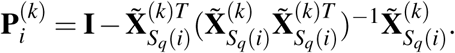 After 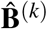 and 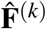 are estimated, the estimate of 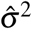 is given by (5). The hyper-parameter ***τ*** in ridge regression (4) or (6) is selected by 5-fold cross-validation.

### 1.4 FSSEM algorithm

In this section, we will develop the FSSEM algorithm to minimize the objective function *J*(**B**, **F**) in (3) with the initial values of **B**^(*k*)^ and **F**^(*k*)^ given in (7). The objective function is non-convex due to the log-determinant term, and non-smooth due to the *ℓ*_1_ norm terms. Recently, the proximal alternating linearized minimization (PALM) method (Bolte et *al.*, 2014) was developed to solve a broad classes of non-convex and non-smooth minimization problems. We next apply the PALM approach to develop the FSSEM algorithm.

Without loss of generality, we define the proximal operator associated with a proper and lower semi-continuous function *h*(**x**): ℝ^*d*^ → (− ∞, +∞] as 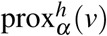
= argmin_*u*∈ℝ^*d*^_ {*h*(*u*) + *α*/2∥*u* − *v*∥^2^}, where *α* > 0 and *ν* ∈ ℝ^*d*^ are given. We also define the fused lasso signal approximator (Friedman et *al.*, 2007; Hoefling, 2010) on *x* = [*x*_1_,*x*_2_] as the following proximal operator:

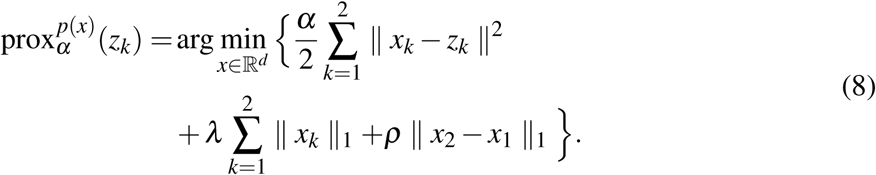

The solution (*x*_1_ (*λ*),*x*_2_(*λ*)) of (8) at *λ* = 0 can be found as

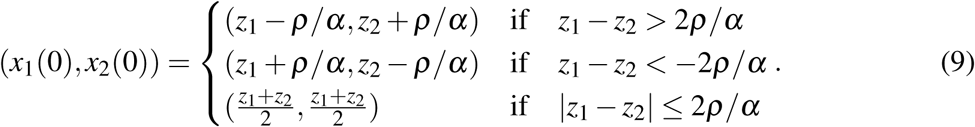

Defining soft-thresholding function *S*(*β*, *λ*) as

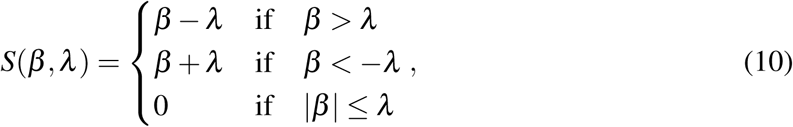

the solution of (8) at *λ* > 0 is given in terms of the soft-thresholding operator as follows (Friedman et al., 2007):

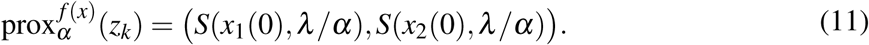

Minimizing (3) w.r.t. **F**^(*k*)^ yields 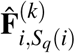
in (7). Substituting 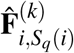
in (7) into (3) gives

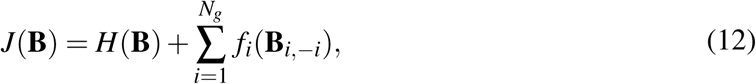

where

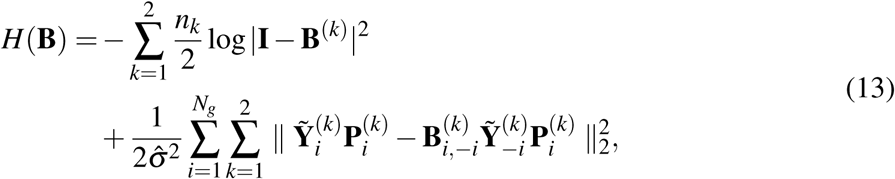

and

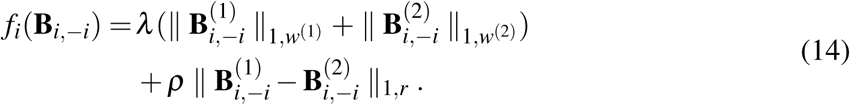

Using the inertial version of the PALM approach (Pock and Sabach, 2016), the FSSEM algorithm efficiently minimizes the non-convex non-smooth function *J*(**B**) with the block coordinate descent (BCD) method in an iterative fashion. More specifically, in each cycle of the iteration, *J*(**B**) is minimized successively w.r.t. 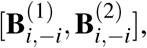
while 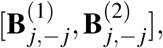
*j* = 1,…, *n*, *j* ≠ *i* are fixed.

Let us consider updating the ith block of variables 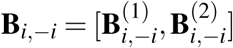
in the (*t* + 1)th cycle. Let **B**[*t*] = [**B**^(1)^[*t*],**B**^(2)^[*t*]] be the estimate of **B** in the *t*th cycle. Define **B̃**_*i*,−*i*_ = **B**_*i*,−*i*_[*t*_1_] + *α_t_*(**B**_*i*,−*i*_[*t*_1_] − **B**_*i*,−*i*_[*t*_1_ − 1]), where *t*_1_ = *t* + 1, ∀*i* < *j*, *t*_1_ = *t*, ∀*i* > *j*, and *α_t_* is a constant in the interval [0,1]. We obtain **B**_*i*,−*i*_ from the FLSA proximal operator (11) as follows:

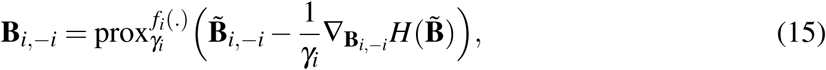

where 1 /*γ_i_* is the step-size for the *i*-th block that will be given later, and ∇_**B**_*i*,−*i*__***H***(**B̃**) is the partial derivative of ***H***(**B**) w.r.t. **B**_*i*,−*i*_ at **B̃**.

Since 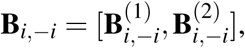
we have 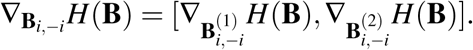
The determinant of **I** − **B**^(*k*)^ can be expressed as 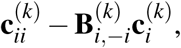
where 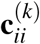
is the (*i*, *i*) co-factor of **I** - **B**^(*k*)^, and the *j*th entry of the (*n* − 1) × 1 column vector 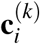
is the co-factor of **I** − **B**^(*k*)^ corresponding to the *j*th entry of 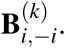
Defining 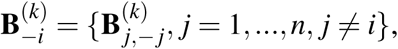
we can write 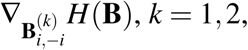
with 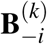
fixed, as follows

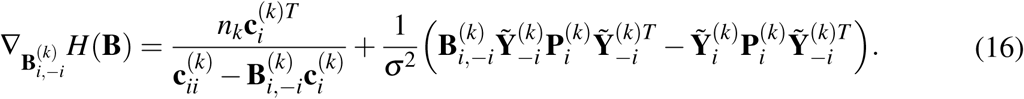

In Supplementary Text S, we prove that given 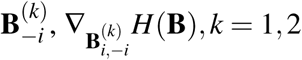
are Lipschitz continuous.

Specifically, we can write 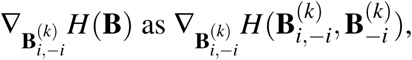
which satisfies:

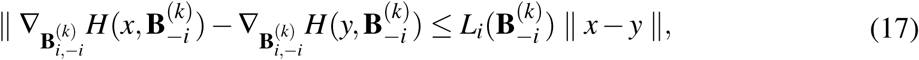

where the Lipschitz constant 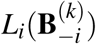
is derived in the Supplementary Text S, and is given by

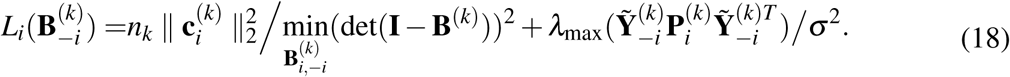

Here 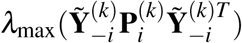
is the maximum eigenvalue of 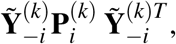
and the value of 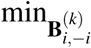
det(**I** − **B**^(*k*)^)^2^ can be computed by solving the optimization problem as shown in (S12) in Supplementary

#### Algorithm 1

Fused Sparse SEM (FSSEM)

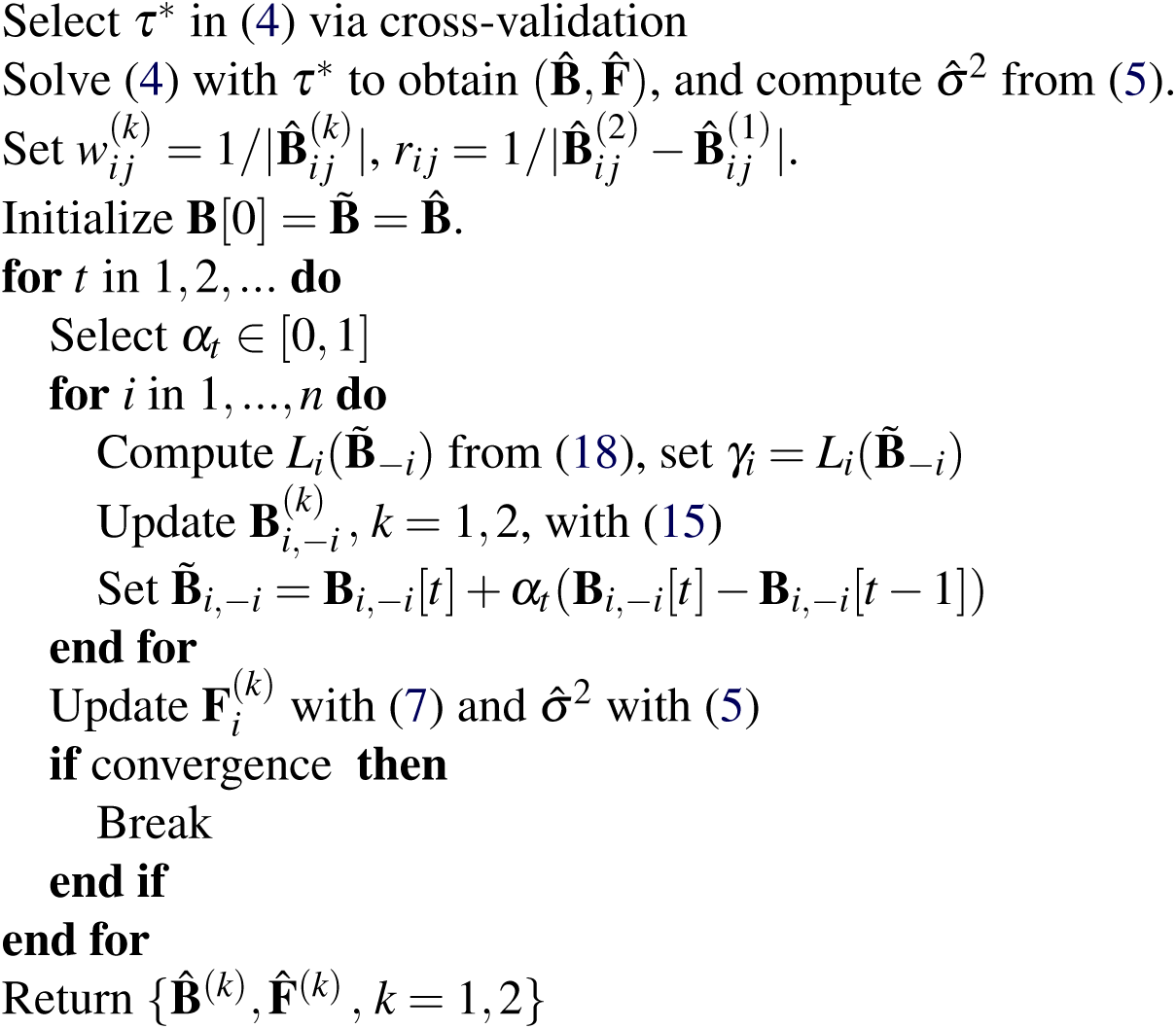

Text S. Let *L_i_*(**B**_−*i*_) = 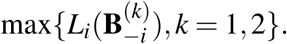
Then, the step size is chosen to be 1/*γ_i_* = 1/*L_i_*(**B̃**_−*i*_.

The FSSEM algorithm is summarized in Algorithm 1. The convergence criterion is defined as

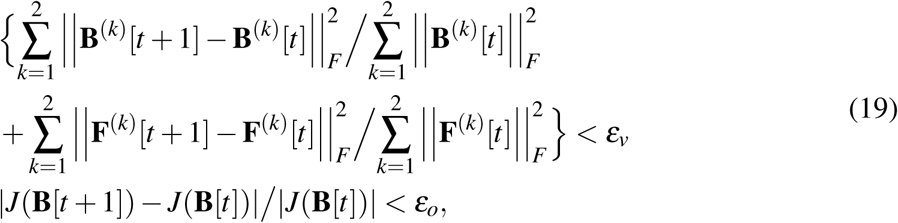

where *ε_ν_* > 0 and *ε_o_* > 0 are pre-specified small constants. Since the objective function is not convex, it is not guaranteed that the FSSEM algorithm converges to the global minimization. However, we prove in Supplementary Text S that the FSSEM algorithm always converges to a stationary point of the objective function. Note that if we drop the fused lasso term *ρ* ∥**B**^(1)^ − **B**^(2)^∥_1,*r*_ in (3), then minimizing *J*(**B**, **F**) is equivalent to estimating two network matrices **B**^(1)^ and **B**^(2)^ separately. The BCD approach used in FSSEM can also be employed to solve this problem, because the proximal operator in (15) can be easily solved in terms of the soft-thresholding function *S*(*β*, *λ*) defined in (10). This BCD approach is much more efficient than the SML algorithm in (Cai *et al.*, 2013), which employs the element-wise coordinate ascent approach. Parameters *λ* and *ρ* in (3) can be determined with cross-validation (CV). In Supplementary Text S, we derive the expression for the maximum values of *λ* and *ρ* and describe the CV process.

## Results

### 2.1 Computer simulations

In this section, we conduct simulation studies to compare the performance of the FSSEM algorithm with that of the SML algorithm (Cai et *al.*, 2013). FSSEM estimates network matrices **B**^(1)^ and **B**^(2)^ jointly, while SML estimates **B**^(1)^ and **B**^(2)^ separately. Other algorithm such as AL-based (Logsdon and Mezey, 2010) and QDG (Neto et *al.*, 2008) algorithms are available to estimate **B**^(1)^ and **B**^(2)^ separately. However, as shown in (Cai et *al.*, 2013), SML algorithm outperforms AL-based and QDG algorithms. Therefore, only SML is considered in performance comparison.

Following the setup of (Cai et *al.*, 2013), both directed acyclic networks (DAG) and directed cyclic networks (DCG) are simulated in our experiments. Specifically, the adjacency matrix **A**^(1)^ of a DAG or DCG of 10 or 30 gene nodes with expected number of edges per gene *d* = 3 is generated for the GRN under condition 1. Another adjacency matrix **A**^(2)^ was generated by randomly change 10% entries of **A**^(1)^, and the probabilities of changes of entries from 0 to 1 and from 1 to 0 are equal. A network matrix **B**^(1)^ was generated from **A**^(1)^ as follow. For any entry 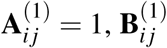
is generated from a random variable uniformly distributed over interval [0.5,1] or [−1, −0.5]; for all 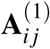
= 0, we set 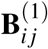
= 0. The second network matrix **B**^(2)^ was generated from **A**^(2)^ and **B**^(1)^ as follow. For all 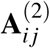
= 0, we set 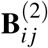
= 0; for all 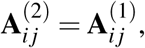
we set 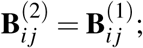
and for all 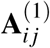
= 0 but 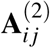
= 1, we generate 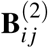
from a random variable uniformly distributed over interval [0.5,1] or [−1, −0.5]. The genotypes of eQTLs were simulated from an F2 cross. Values 1 and 3 were assigned to two homozygous genotypes, respectively, and value 2 to the heterozygous genotype. Then, **X**^(1)^ and **X**^(2)^ were generated from ternary random variables taking on values {1,2,3} with corresponding probabilities {0.25,0.5,0.25}. The number of eQTLs per gene *n_e_* was chosen to be either 1 or 3, and effect sizes of all eQTLs were set to 1 in **F**^(1)^ and **F**^(2)^. Error terms **E**^(1)^ and **E**^(2)^ were independently sampled from Gaussian random variables with zero mean and variable **σ**^2^; *μ*^(1)^ and *μ*^(2)^ were set to zero vectors; and the sample sizes *n*_1_ and *n*_2_ vary from 100 to 1,000. Finally, **Y**^(*k*)^ was calculated as **Y**^(*k*)^ = (**I** − **B**^(*k*)^)^−1^ (**F**^(*k*)^**X**^(*k*)^ + **E**^(*k*)^), where *k* = 1,2.

For each configuration of the two GRNs, 30 replicates of the GRN were simulated. For each replicate, SML and FSSEM were used to infer network matrices **B**^(1)^ and **B**^(2)^. Hyper parameter of SML and FSSEM algorithms were determined with 5-fold CV. Power of detection (PD) and false discovery rate (FDR) for detecting network edges were calculated from **B**^(1)^ and **B**^(2)^ estimated from the data of each of 30 network replicates. The differential network was defined as Δ**B** = **B**^(2)^ − **B**^(1)^, and PD and FDR for the differential network were calculated accordingly.

The results for DAGs with *n* = 30, *n_e_* = 3 and **σ**^2^ = 0.25 are depicted in Figure 1, and results of DAGs under other settings are given in Figures S1 and S2 in Supplementary Text S. First, let us look at the PD and FDR of **B**^(1)^ and **B**^(2)^ in the left panel of Figure 1. FSSEM offers almost the same PD as SML, but slightly lower FDR when σ^2^ = 0.25. As shown in Figures S1 and S2, when **σ**^2^ = 0.01, FSSEM and SML have almost same PD and FDR, and increasing *n_e_* from 1 to 3 slightly reduced FDR for both FSSEM and SML. Overall, FSSEM and SML have similar performance for the estimates of **B**^(1)^ and **B**^(2)^. Next, let us look at the PD and FDR of **B**^(2)^ − **B**^(1)^ in the right panel of Figure 1. FSSEM exhibits almost the same PD as SML, but it offers much smaller FDR than SML. Specially, the FDR of FSSEM is < 0.2, but the FDR of SML > 0.8. Similar trends are seen in Figure S1 and S2 for all network settings considered. Again, increasing the number of eQTL *n_e_* from 1 to 3 slightly reduces the FDR. For the GRNs of *n* = 30 genes, there are 2(*n*^2^ − *n*) = 1740 unknown entries in **B**^(1)^ and **B**^(2)^ to be estimated, and there are 2*nn_e_* unknown entries in **F**^(1)^ and **F**^(2)^. Interestingly, the performance of both FSSEM and SML did not change much, when the sample size *n*_1_ + *n*_2_ varied from 400 to 4,000.

**Figure 1.**
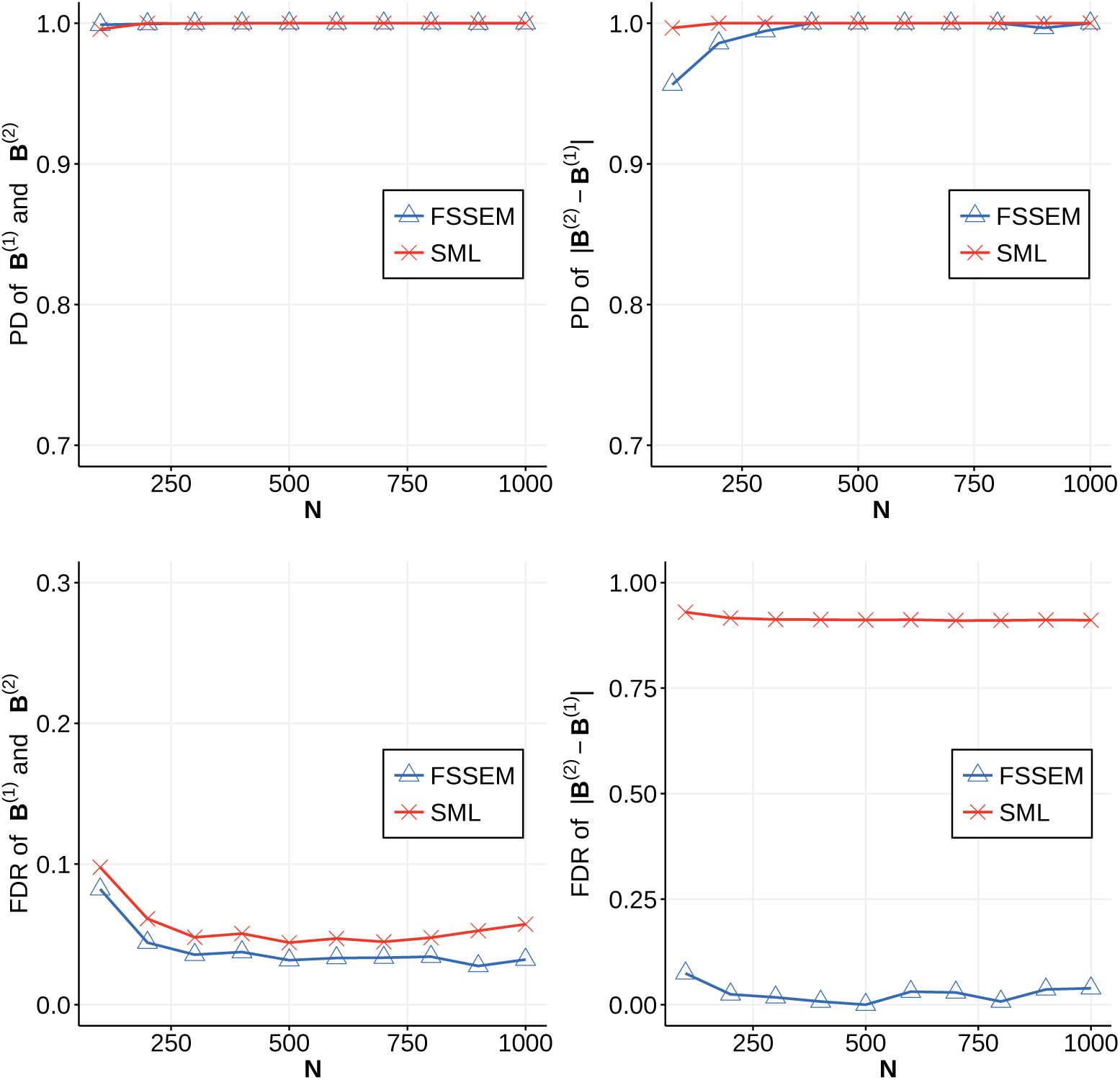
Performance of FSSEM and SML for the DAG with *n* = 30 genes and *n_e_* = 3 eQTLs per gene. The number of samples *n*_1_ = *n*_2_ varies from 100 to 1,000 and noise variance σ^2^ = 0.25. PD and FDR were obtained from 30 network replicates.

Simulation results for DCGs with *n* = 30, *n_e_* = 3 and σ^2^ = 0.25 are depicted in Figure 2, and results of DCGs under other settings are shown in Figures S3 and S4 in Supplementary Text S. As shown in the left panel of Figure 2, FSSEM offers slightly better PD and FDR for the estimates of **B**^(1)^ and **B**^(2)^ than SML. The results for ΔB shown in the right panel indicate that FSSEM offers similar PD comparing with SML, but it exhibits much smaller FDR, as also observed in Figure 1 for DAG networks. Similar trends are also observed for other network settings in Figures S3 and S4. Clearly, FSSEM outperforms SML consistently in terms of both PD and FDR for both DAG and DCG networks. For the convenience of comparison, the simulation results of DAG and DCG with *n*_1_ = *n*_2_ = 500, *n_e_* = 3 and σ^2^ = 0.25 are summarized in Table 1, which clearly shows that FSSEM outperforms SML. Particularly, FSSEM offers much lower FDR than SML in estimating ΔB, the matrix of the differential GRN.

**Figure 2.**
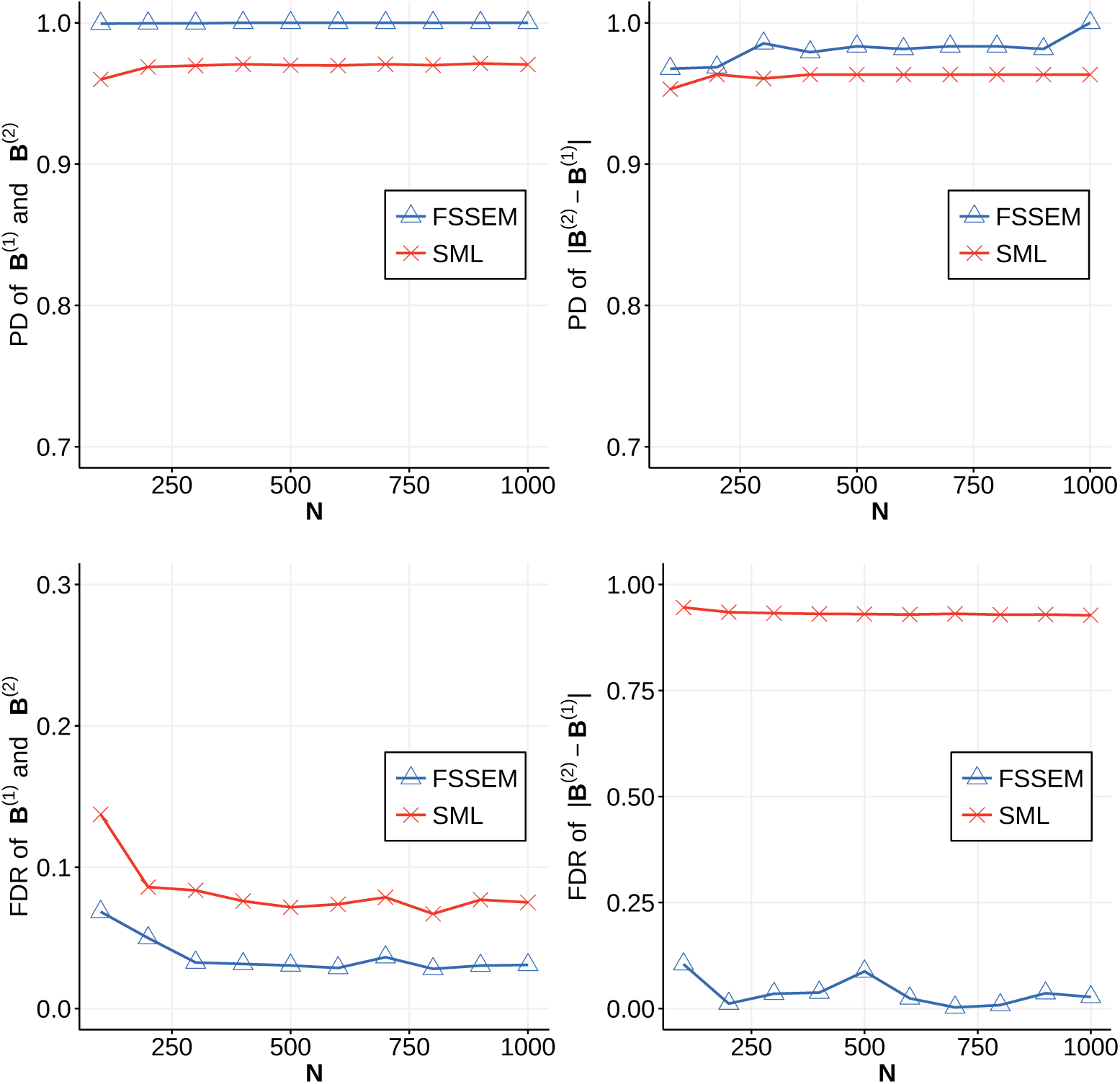
Performance of FSSEM and SML for the DCG with *n* = 30 genes and *n_e_* = 3 eQTLs per gene. The number of samples *n*_1_ = *n*_2_ varies from 100 to 1,000 and noise variance σ^2^ = 0.25. PD and FDR were obtained from 30 network replicates.

**Table 1.** Performance of FSSEM and SML algorithms. Expected number of eQTLs per gene is *n_e_* = 3 and noise variance σ^2^ = 0.25. PD and FDR were obtained from 30 network replicates.

| Network | $n$ | FSSEM | | | | SML | | | |
| --- | --- | --- | --- | --- | --- | --- | --- | --- | --- |
| | | $PD_B$ | $FDR_B$ | $PD_{\Delta B}$ | $FDR_{\Delta B}$ | $PD_B$ | $FDR_B$ | $PD_{\Delta B}$ | $FDR_{\Delta B}$ |
| DAG | 10 | 1.000 | 0.037 | 1.000 | 0.000 | 1.000 | 0.054 | 1.000 | 0.889 |
|  | 30 | 1.000 | 0.032 | 1.000 | 0.000 | 1.000 | 0.044 | 1.000 | 0.911 |
| DCG | 10 | 1.000 | 0.028 | 1.000 | 0.016 | 0.879 | 0.082 | 0.856 | 0.912 |
|  | 30 | 1.000 | 0.004 | 1.000 | 0.057 | 0.962 | 0.083 | 0.955 | 0.920 |

### 2.2 Real data analysis

In (Lu *et al.*, 2011), gene expression levels in 42 tumors and their adjacent normal tissues of non-smoking female patients with lung adenocarcinoma were measured with 54,675 probe sets from Affymetrix Human Genome U133 Plus 2.0 Array. The genotypes of single nucleotide polymorphisms (SNPs) in the same set of tissues were obtained using 906,551 SNP probes from Affymetrix Genome-Wide Human SNP 6.0 array. We applied FSSEM to this data set to infer GRNs in lung cancer and normal tissues.

Both gene expression and SNP data in the gene expression omnibus database (GSE33356) were downloaded. The R package affy (Gautier *et al.*, 2004) was employed to transform raw micro-array data to normalized gene expression levels. Specifically, the raw gene expression data in the custom CDF format (Dai *et al.*, 2005) were normalized using the robust multi-array average (RMA) method (Bolstad *et al.*, 2003; Irizarry *et al.*, 2003a,b). In total, gene expression levels of 18,807 genes with their Entrez IDs were obtained from 54,675 probe sets. the genotypes of the 906,551 SNP probes in the 84 tissue samples were transformed to values {0,1,2} using the following mapping: AA → 0, AB → 1 and BB → 2. The missing genotypes of SNP probes were imputed by randomly sampling from {0,1,2} using the observed probabilities of genotypes of each SNP. Finally, R package MatrixEQTL (Shabalin, 2012) was adopted to identify *cis*-eQTLs of genes. In total, 1,456 genes were found to have at least one *cis*-eQTLs within 10^6^ base pairs from the open reading frame (ORF) of the gene at an FDR = 0.01.

Since the number of samples available is 84, which may be too small to be used to reliably infer the network of 1,456 genes with eQTLs, we selected a subset of the 1,456 genes as follow. In (Greene *et al.*, 2015), gene interactions in 144 human tissues and cell types were inferred by integrating a collection of data sets covering thousands of experiments reported in more than 14,000 distinct publications. Each identified interaction between a pair of genes was assigned a confidence score or posterior probability in the Bayesian data integration process. The tissue-specific gene networks are all available in the GIANT (http://hb.flatironinstitute.org) database. For each pair of genes among the 1,456 genes with eQTLs, we searched the GIANT database to see if they interact with each other in the lung network constructed from the gene expression data. We identified the following 19 genes that interact with at least one another gene with high confidence (posterior probability ≥ 0.80): UBA2, CCT7, COX6B1, DBI, DKC1, ETFA, NACA, PSMC4, RPS6, SNRPF, BRIX1, KARS, ECHS1, ATP5G3, UBE2N, CDC123, VBP1, PSMD10, and BTF3. We extracted interactions among these 19 genes with posterior probability ≥ 0.80, and refer to this sub-network as the GIANT reference network. Since the dataset GSE33356 was not used to construct the GIANT network, we would compare the GRN inferred from the GSE33356 by our FSSEM algorithm with the GIANT network.

We applied the FSSEM algorithm to the 84 samples of expression levels of the 19 selected genes and the genotypes of their eQTLs to infer the networks of these genes in lung cancer and normal tissue. An edge from gene *j* to gene *i* was detected if 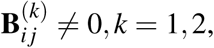 where **B**^(1)^ and **B**^(2)^ specify the networks in normal and tumor tissues, respectively. FSSEM yielded a network matrix **B**^(1)^ with 93 nonzero entries, or a network with 93 edges, and these edges were regarded as significant gene interactions in normal tissues. This network referred to as the FSSEM network was compared with the GIANT reference network. It was found that 76.9% edges in the FSSEM network **B**^(1)^ were also in the GIANT network. This shows that the FSSEM network identified from a small independent samples is in good agreement with the GIANT reference network identified from a large number of data samples.

We also identified the differential network based on Δ**B** = **B**^(2)^ – **B**^(1)^. Since small changes of coefficients **B**_*ij*_ may not have much biological effect, we regarded the regulatory effect of gene *j* and *i* to be different if 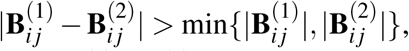 which ensured that there is at least one-fold change relative to min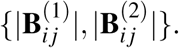 However, when one of 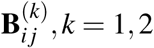 is zero or near zero, this criterion still fails to filter out very small changes Δ**B**. To avoid this issue, we added another criterion. Specifically, we obtained all nonzero entries of **B**^(*k*)^, *k* = 1,2, and compute the 20 percentile value of all nonzero 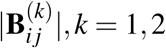 as *η*. Then, we defined the second criterion as max 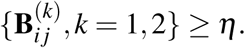. We employed the stability selection technique (Meinshausen and Bühlmann, 2010) to identify the differential network reliably. Specifically, we used 5-fold cross-validation to determine optimal values of *λ* and *ρ*, denoted as *λ*\* and *ρ*\*, respectively. A set of 21 samples are randomly selected from 42 cancer samples, and another set of 21 corresponding samples were selected from 42 normal samples. This data set of 42 samples was used by FSSEM algorithm to infer **B**^(1)^ and **B**^(2)^ with *λ* = *λ*\*and *ρ* = *ρ*\*. The changed edges were identified based on Δ**B** and the two criteria described earlier. This process of random sampling and network inference was repeated 100 times, and a changed edge was declared to be significant, if it was detected more than 80 times.

Stability selection yielded 16 edges that changed significantly. The differential network that is formed by these 16 edges is shown in Figure 3. Most edges in the differential network are connected with genes PSMD10, RPS6, BRIX1, VBP1 and SNRPF. As will be discussed in the next section, these five genes have been reported in a number of experimental results to be implicated in lung and other cancers.

**Figure 3.**
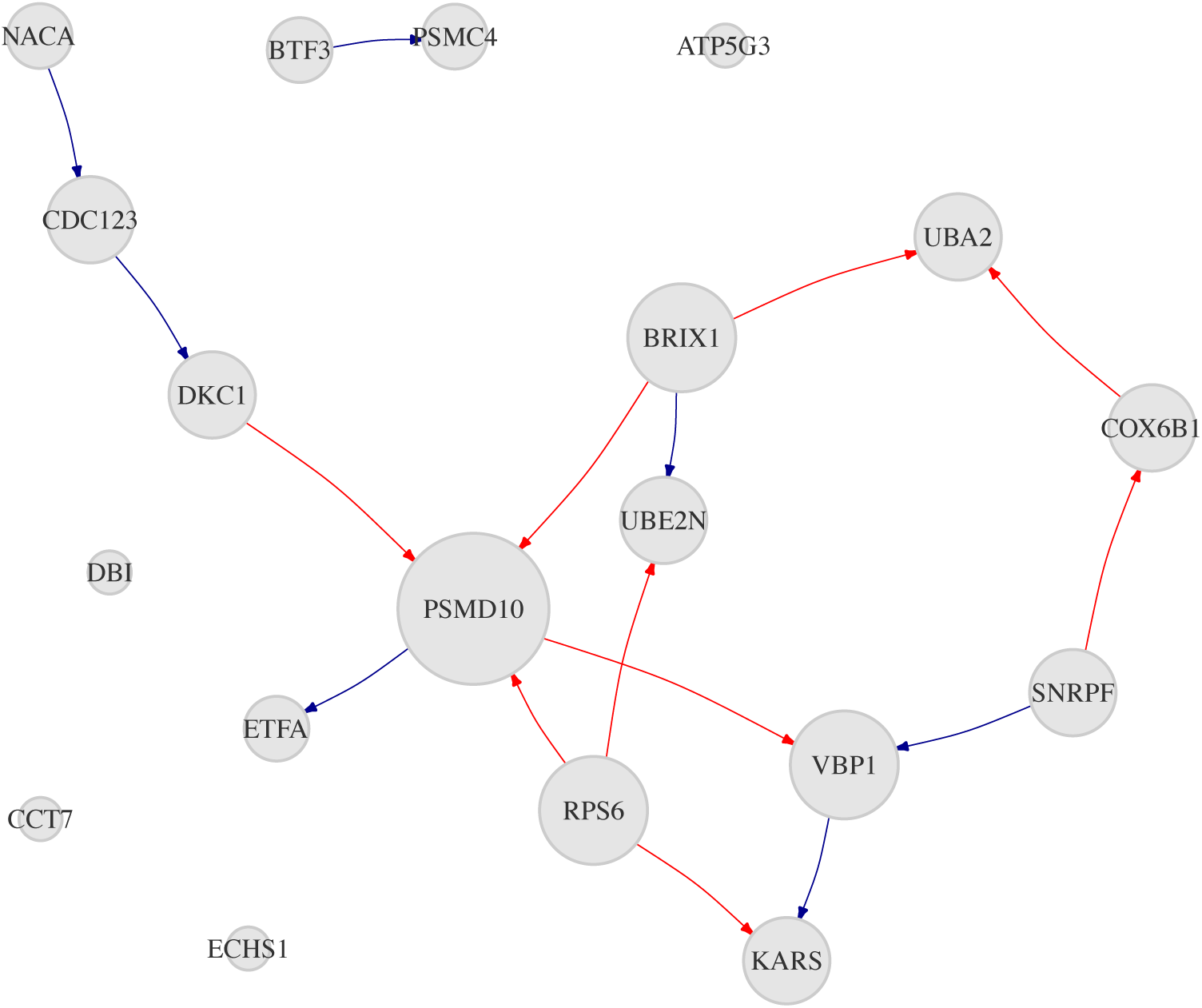
The differential regulatory network of 19 genes inferred from gene expression and eQTL data with the FSSEM algorithm. The size of a node is proportional to its degrees.

## Discussion

In this paper, we developed a very efficient algorithm, named FSSEM, for joint inference of two similar GRNs by integrating genetic perturbations with gene expression data under two different conditions with the structural equation model. An R package implementing the FSSEM algorithm is available, which provides a useful tool for inferring GRNs. Computer simulations showed that our FSSEM offered much better accuracy in identifying changed gene interactions than the approach that infers two GRNs separately. Particularly, the FDR of gene interactions in the differential GRN estimated by FSSEM was significantly lower than that resulted from the method estimating two GRNs separately. This result is expected because FSSEM exploits the similarity in the two GRNs and penalizes the changes of gene interactions in the inference process.

Analysis of a data set of lung cancer and normal tissues with FSSEM detected most gene interactions identified in another study that exploited a large number of data sets. Real data analysis also identified several genes that may be involved in cancer development. Specifically, PSMD10 is aberrantly expressed in various cancers, and its expression level is inverse correlated with the expression level of miR-605 (Li *et al.*, 2014), which is repoted to be associated with lung cancer (Yin *et al.*, 2016). RPS6 is a component of the 40S ribosomal subunit; its expression has been shown to increase significantly in non-small cell lung cancer (NSCLC) (Chen *et al.*, 2015). Additionally, RPS6 is regulated in multiple signal pathways, such as the Akt2/mTOR/p70S6K signaling pathway (Yano *et al.*, 2014), that are closely related to the progression of NSCLC (Chen *et al.*, 2015). BRIX1 was identified as a key transcription factor associated with lung squamous cell carcinoma (Zhang *et al.*, 2017) and was also recognized as a key gene in the gastric cancer network (Kutmon *et al.*, 2015). SNRPF is a gene related to mRNA splicing pathway, and it was identified as a biomarker of ovarian cancer (Bengtsson *et al.*, 2007). Recently, it was reported that SNRPF was required for cells to tolerate oncogenic MYC hyperactivation (Hsu *et al.*, 2015). VBP1 is a gene identified to bind to the tumor suppressor gene VHL (Tsuchiya *et al.*, 1996), and it was reported that VBP1 repressed cancer metastasis by enhancing HIF1α degradation induced by pVHL (Kim *et al.*, 2018).

## Supplementary Text S

### Hyper-parameter selection

We use *K*-fold cross-validation (CV) to determine the value of *τ* for ridge regression (4) and values of *λ* and *ρ* for FSSEM, where *K* typically equals to 5 or 10. We search *τ* over a sequence of 50 values increasing from 10^−6^ to 10^2^ evenly on the logarithm scale, and the optimal value of *τ* is chosen to minimize the predication error calculated from the test data. We employ a grid search strategy to determine the optimal values of *λ* and *ρ*. We first determine the maximum value of *λ*, namely *λ*_max_, then choose a set of *k*_1_, values for *λ*, denoted as sequence
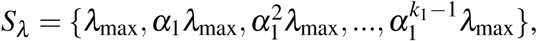 where 0 < *α*_1_ < 1. For each value of *λ* ∈ *S_λ_*, we find the maximum value of *ρ*, namely *ρ*_max_(*λ*), and then choose a set of *k*_2_ values for *ρ*, denoted as
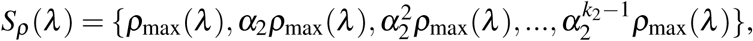 where 0 < *α*_2_ < 1. This gives a set of *K* = *k*_1_*k*_2_ pairs of (*λ*, *ρ*), and CV is carried out over this parameter space. The optimal values of *λ* and *ρ* are chosen to minimize the likelihood calculated from the test data.

Next, we derive the maximum values of *λ* and *ρ* needed in CV. The value *λ*_max_ yields **B**^(1)^ = **B**^(2)^ = **0**, and can be found from the result in (Cai *et al.*, 2013) as follows:

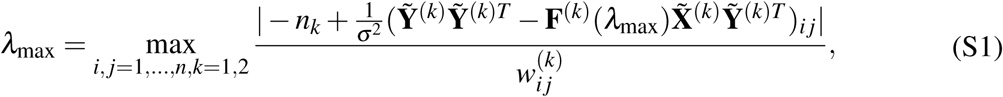

where **F**^(*k*)^(*λ*_max_) can be determined from (7) by setting
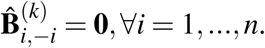 When *ρ* = *ρ*_max_(*λ*), minimizing *J*(**B**, **F**) in (3) yields **B**^(1)^ = **B**^(2)^. Therefore, we let **B**^(1)^ = **B**^(2)^ = **B**, which yields

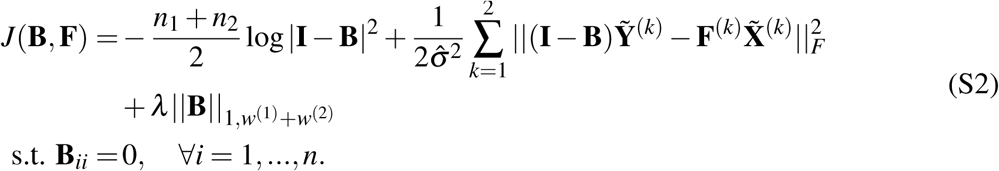

Then, we use the SML algorithm (Cai *et al.*, 2013) or the BCD approach of FSSEM mentioned in the main text to get **B̂** that minimizes ***J***(**B**, **F**) in (S2). Using the fact that when *ρ* = *ρ*_max_(*λ*), the sub-gradient of ***J***(**B**, **F**) in (3) w.r.t. **B̂**_*i*,*j*_ equals to zero at **B̂**^(1)^ = **B̂**^(2)^ = **B̂**, we obtain

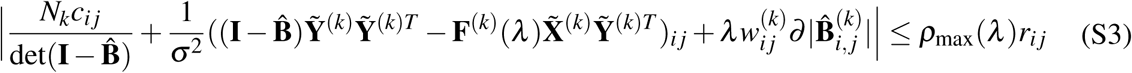

where *∂*(|*β*|) is the sub-gradient of |*β*|, and *∂*(|*β*|) = 1, if *β* > 0, *∂*(|*β*|) = –1, if *β* < 0, and *∂*(|*β*|) ∈ [–1, 1] if *β* = 0, and *c_ij_* is the (*i*, *j*) co-factor of **I** – **B̂**. From (S3), we obtain

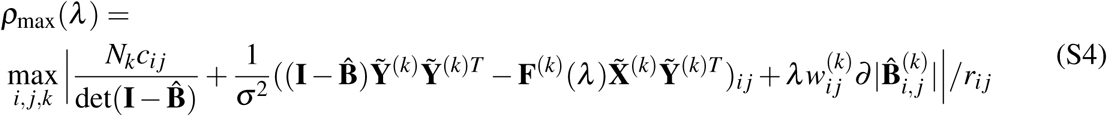

### Convergence analysis

When the objective function in an optimization problem is non-convex and non-smooth, it is possible that the coordinate descent method fails to converge. We next prove that the FSSEM algorithm converges to a stationary point, because the objective function satisfies the conditions for the convergence of the PALM method specified in (Bolte et al., 2014). Specifically, *J*(**B**) in (15) has the following properties:

1. inf *H* (**B**) > –∞ and inf *f_i_*(**B**_*i*,–*i*_) > –∞, *i* = 1,…*n*.
2. ∇_**B**_*i*, –*i*__*H*(**B**),*i* = 1,…,*n*, is gradient Lipschitz continuous with constant *L_i_*(**B**_–*i*_) when **B** ∈ dom*H* = {**B**: det(**I** – **B**^(*k*)^) ≠ 0,*k* = 1,2}:

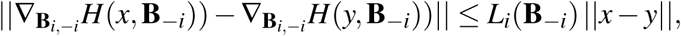
3. *H*(**B**) has continuous first and second derivatives when **B** ∈ dom*J* = {**B**: det(**I** – **B**^(*k*)^) ≠ 0,*k* = 1,2}.
4. *J*(**B**) satisfies the Kurdyka-Łojasiewicz(KL) property.

Note that properties 1-3 are identical to the properties in assumption B of (Bolte et *al.*, 2014), and these 4 properties guarantee that FSSEM algorithm converges to a critical point. First, it is apparent that *H*(**B**) > –∞ and therefore *J*(**B**) > –∞, ∀**B** ∈ dom*J*. Second, it is not difficult to show that *H*(**B**) is differentiable w.r.t. **B**_*i*, –*i*_, *i* = 1,…,*n* and the first-order and second-order derivatives are continuous in dom*H*. Therefore, property 3 is satisfied.

Third, we prove in the next section that *H*(**B**_*i*,–*i*_,**B**_–*i*_) is gradient Lipschitz continuous with constant *L_i_*(**B**_–*i*_) given in (21). Moreover, based on assumption B(iii) of (Bolte *et al.*, 2014), *L_i_*(**B**_–*i*_) guarantees that proximal steps in the FSSEM algorithm remain well-defined, because we have

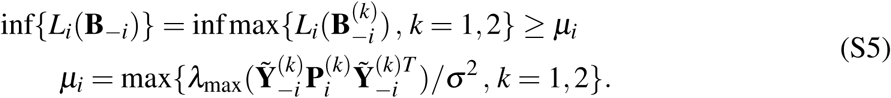

Finally, we prove that property 4 holds. The non-smooth functions *f_i_*(**B**_*i*,–*i*_) in *J*(**B**) is the sparse fused lasso penalty term w.r.t. **B**_*i*,–*i*_, and it is semi-algebraic as shown in (Xu and Yin, 2013). The *ℓ*_2_ norm
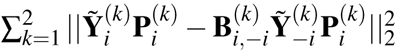 is apparently semi-algebraic. We next prove that
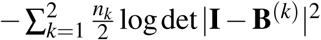 is semi-algebraic too. We can regard
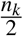 log det |**I** – **B**^(*k*)^|^2^, *k* = 1,2, as a composite function of **B**^(*k*)^ as follows

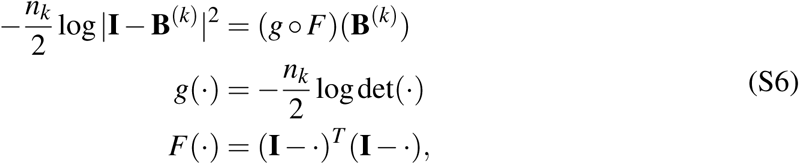

Function *g*(⋅) is locally convex function (Boyd and Vandenberghe, 2004). Based on the result in (Xu and Yin, 2013), it is not difficult to show that *g*(⋅) satisfies the KL property, and it can be shown that function *F*: ℝ^*n*×*n*^ → ℝ^*n*×*n*^ is continuously differentiable in dom *J*. As all terms of *J*(**B**) are KL functions, the sum of theses KL functions should satisfy the KL property (Li and Pong, 2017). This completes the proof that *J*(**B**) satisfies properties 1-4.

### Derivation of the Lipschitz constant of ∇_**B**_*i*,–*i*__*H*(**B**)

In this section, we derive the Lipschitz constant of ∇_**B**_*i*,–*i*__*H*(**B**) in (16), where we drop index *t* in *B*[*t*] for notational simplicity. From (16), we obtain the following:

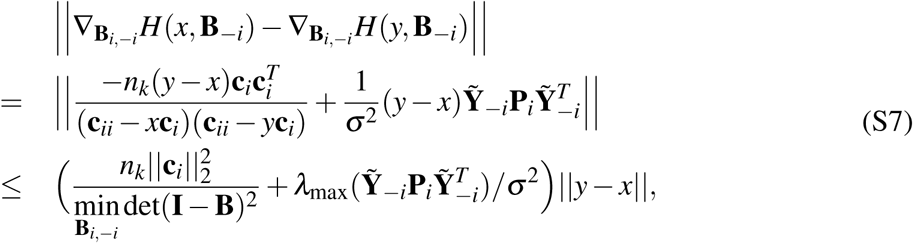

where min_**B**_*i*,–*i*__ det(**I** – **B**) is the minimal value of det(**I** – **B**) for a given **B**_–*i*_ ∈ dom*J*, and *λ*_max_
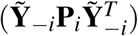 is the maximum eigenvalue of
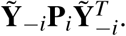 The Lipschitz constant of ∇_**B**_*i*,–*i*__*H*(**B**) is then given by

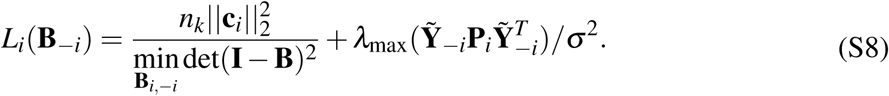

The value of min_**B**_*i*,–*i*__ det(**I** – **B**)^2^ can be determined as follows.

Define Θ = **I** – **B**, and let
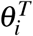 be the *i*th row of Θ, and Θ_–*i*_ be the sub-matrix of Θ that excludes
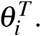 Then, we have,

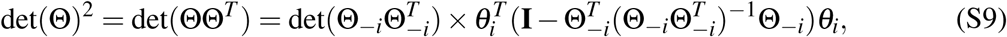

where Θ_−*i*−*i*_ is the submatrix of Θ excluding the *i*th row and the *i*th column. Here we assume **B** ∈ dom *J*, and thus det(ΘΘ^*T*^) > 0. Since 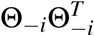
is a submatrix of ΘΘ^*T*^, the Cauchy’s interlacing theorem (Hwang, 2004) implies det 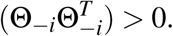
Therefore, 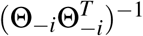
in (S9) exists.

For notational simplicity, we let 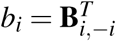
and write (S9) as

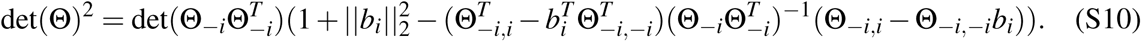

Minimizing det(Θ)^2^ in (S10) w.r.t. *b_i_* yields

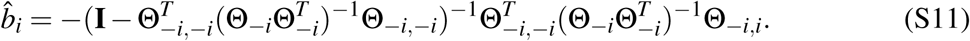

Substituting *b̂_i_* into (S10) gives the minimum value of det(Θ)^2^. In practice, to ensure numerical stability, we modify the *b̂_i_* in (S11) as follows,

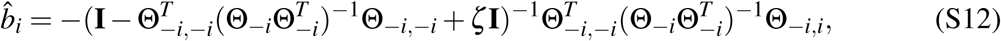

where *ζ* is a small positive constant. This modification can be regarded as minimizing (det(Θ))^2^ in (S11) subject to the constraint ∥*b_i_*∥^2^ ≤ *c*, where *c* is a positive constant. This is a reasonable assumption because in real GRNs, entries of **B**^(*k*)^ are bounded. In our implementation, we chose *ζ* = 10^−16^ and we did not observe any numerical instability in all of our numerical experiments.

**Figure S1.**
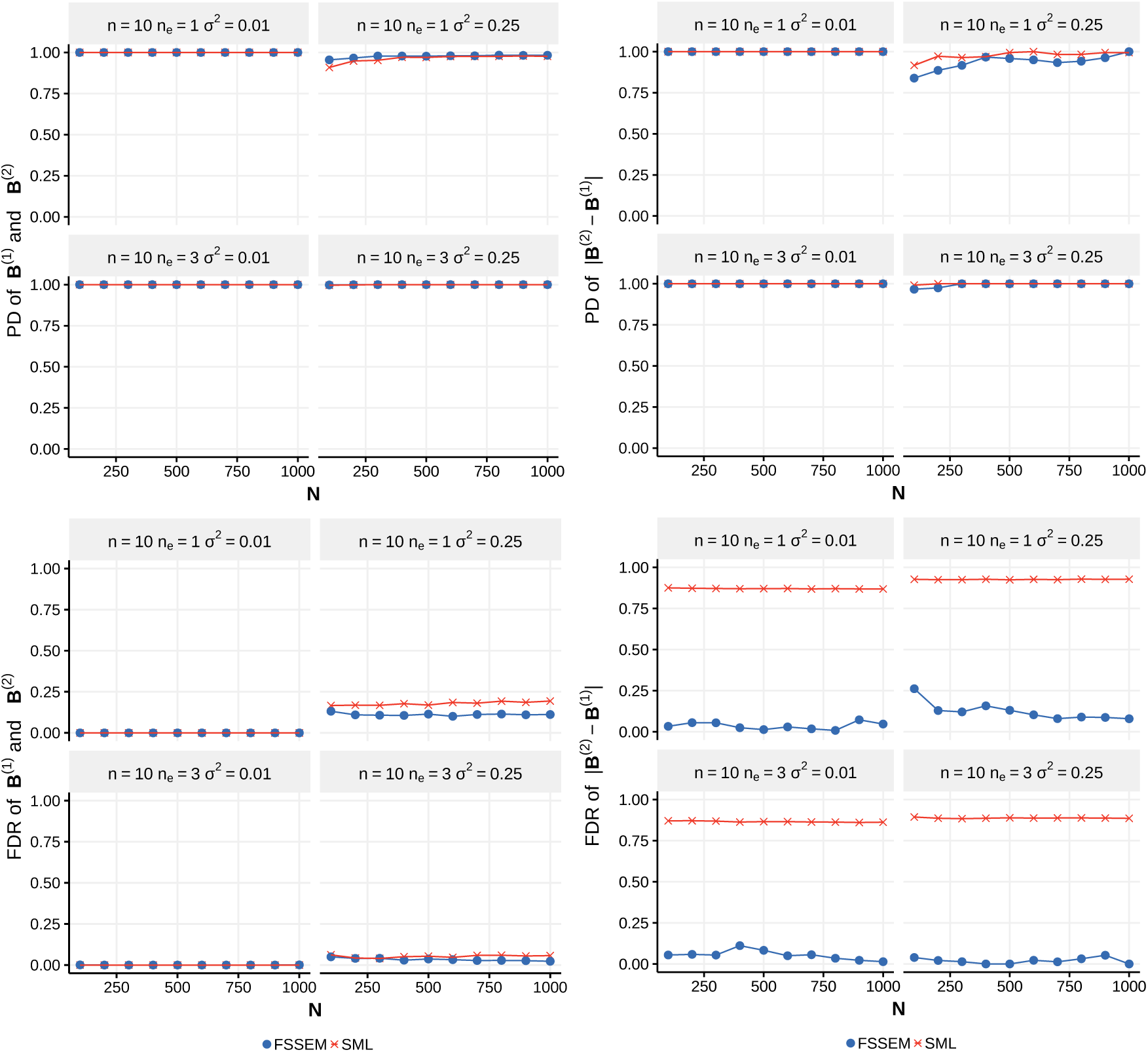
Performance of FSSEM and SML for the DAG with *n* = 10 genes. The number of samples *n*_1_ = *n*_2_ varies from 100 to 1,000 and noise variance σ^2^ = 0.01,0.25. PD and FDR were obtained from 30 network replicates.

**Figure S2.**
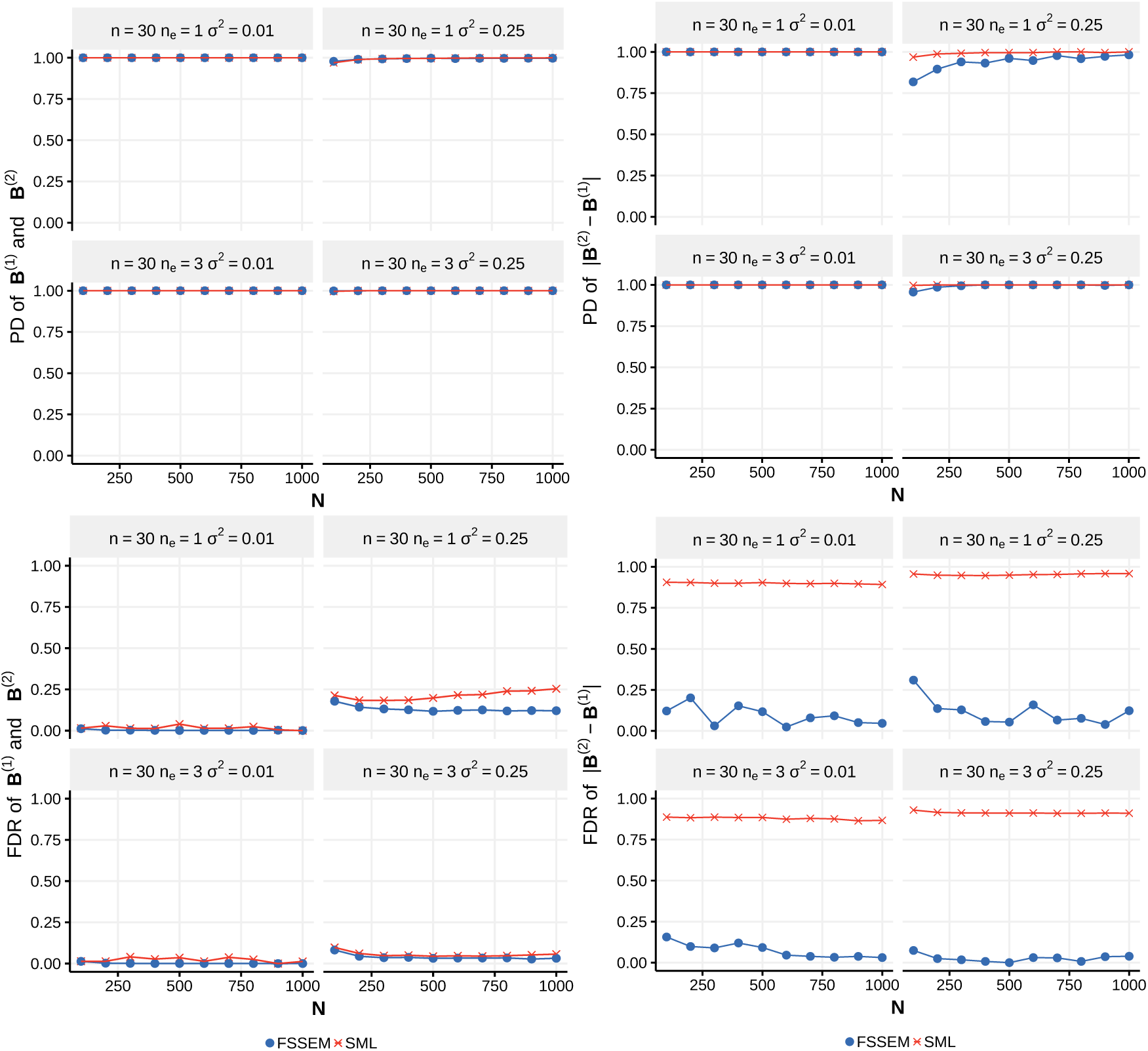
Performance of FSSEM and SML for the DAG with *n* = 30 genes. The number of samples *n*_1_ = *n*_2_ varies from 100 to 1,000 and noise variance σ^2^ = 0.01,0.25. PD and FDR were obtained from 30 network replicates.

**Figure S3.**
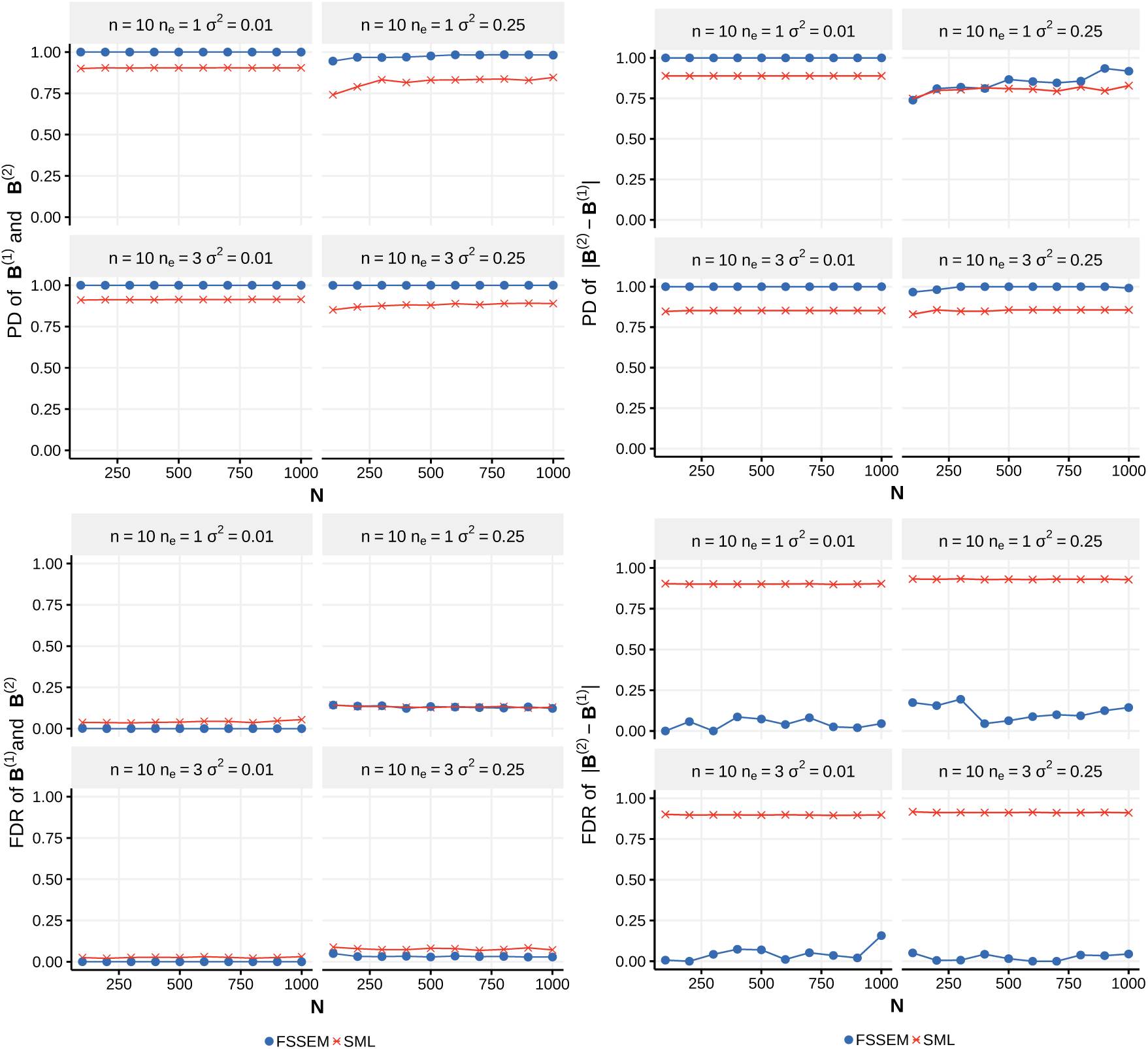
Performance of FSSEM and SML for the DCG with *n* = 10 genes. The number of samples *n*_1_ = *n*_2_ varies from 100 to 1,000 and noise variance σ^2^ = 0.01,0.25. PD and FDR were obtained from 30 network replicates.

**Figure S4.**
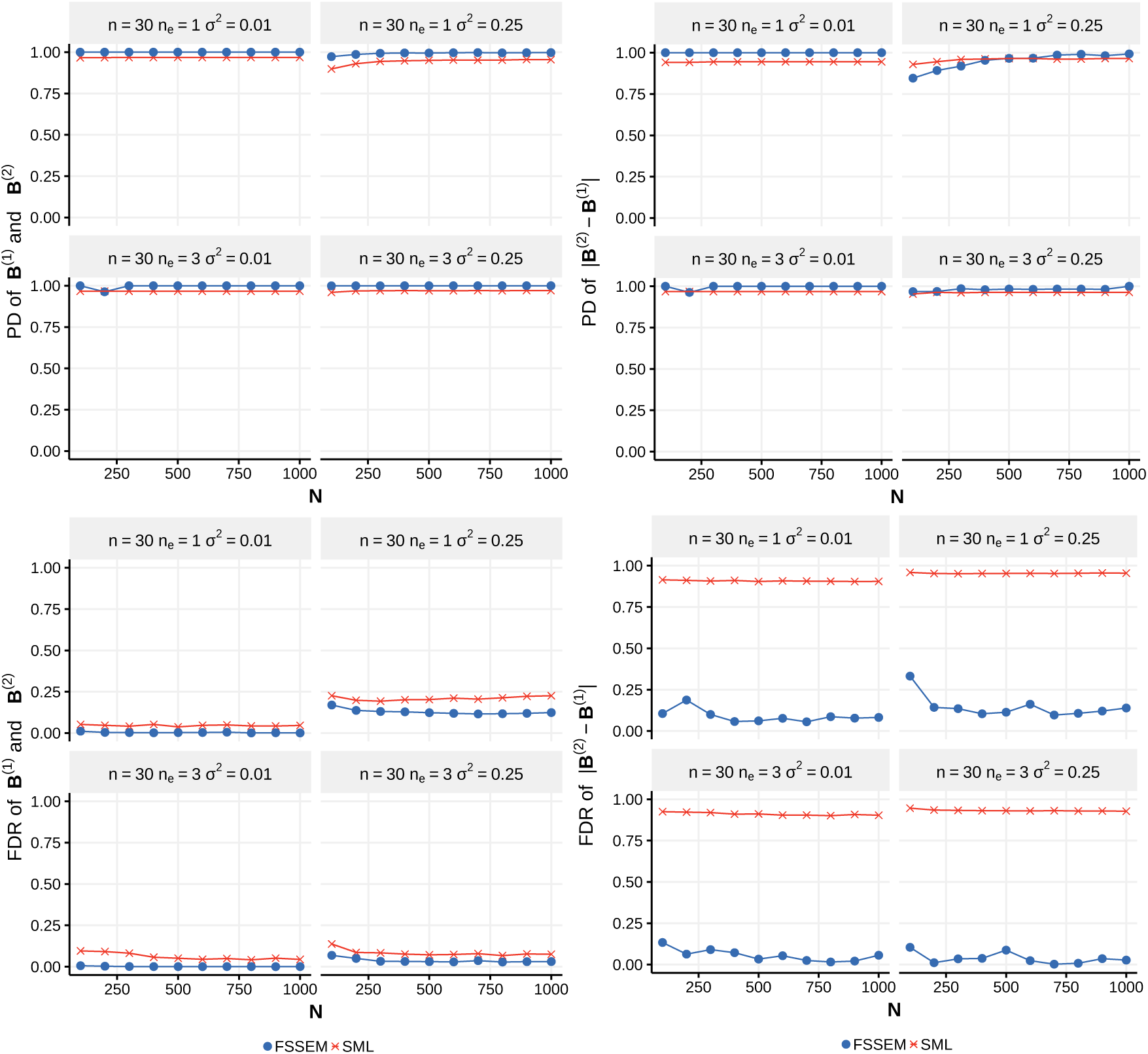
Performance of FSSEM and SML for the DCG with *n* = 30 genes. The number of samples *n*_1_ = *n*_2_ varies from 100 to 1,000 and noise variance σ^2^ = 0.01,0.25. PD and FDR were obtained from 30 network replicates.

